# phasebook: haplotype-aware de novo assembly of diploid genomes from long reads

**DOI:** 10.1101/2021.07.02.450883

**Authors:** Xiao Luo, Xiongbin Kang, Alexander Schönhuth

**Affiliations:** Life Science & Health, Centrum Wiskunde & Informatica, Amsterdam, The Netherlands; Genome Data Science, Faculty of Technology, Bielefeld University, Bielefeld, Germany

**Author notes:** These authors contributed equally to the work (Xiao Luo, Xiongbin Kang).

**Keywords:** Genome assembly, Haplotype, Diploid, Long reads

## Abstract

Haplotype-aware diploid genome assembly is crucial in genomics, precision medicine, and many other disciplines. Long-read sequencing technologies have greatly improved genome assembly thanks to advantages of read length. However, current long-read assemblers usually introduce disturbing biases or fail to capture the haplotype diversity of the diploid genome. Here, we present phasebook, a novel approach for reconstructing the haplotypes of diploid genomes from long reads *de novo*.

Benchmarking experiments demonstrate that our method outperforms other approaches in terms of haplotype coverage by large margins, while preserving competitive performance or even achieving advantages in terms of all other aspects relevant for genome assembly.

## Background

There are generally multiple copies of hereditary material in most eukaryotic organisms, where each copy is inherited from one of the ancestors. The copy-specific nucleic acid sequences are called *haplotypes*, and the haplotype-specific contigs that assembly programs compute for reconstructing them are often referred to as *haplotigs*. Most vertebrates (such as human and mouse) and many higher plants (such as maize and arabidopsis) are diploid, which means that there are two copies for each chromosome.

Haplotype reconstruction plays a crucial role in various disciplines. For example, haplotype information is important in functional genomics since there is widespread allele-specific gene expression across the human genome Tewhey *et al*. (2011); haplotype information also crucially supports studies on population demography, gene flow, and selection in conservation genomics Leitwein *et al*. (2020); the haplotype-specific combinations of genetic variants usually affect disease phenotypes and clinical responses, which is of great concern in precision medicine Muers (2011); Glusman *et al*. (2014).

Over the last few years, long-read sequencing technologies such as single-molecule real-time (SMRT) sequencing of Pacific Biosciences (PacBio) and nanopore sequencing of Oxford Nanopore Technologies (ONT), have improved genome assembly greatly because sequencing reads are sufficiently long (generally ranging from several Kbp to hundreds of Kbp, or even to a few Mbp) to also span more complex repeat regions Jain *et al*. (2018); Wenger *et al*. (2019); Miga *et al*. (2019); Jung et al. (2019).

Although long reads have tremendous advantages in terms of read length in comparison to next generation sequencing (NGS) derived short reads, they suffer from (substantially) elevated sequencing error rates. In particular, PacBio CLR and ONT reads, as the most representative examples of long reads, have sequencing error rate as high as 5 to 15%. While PacBio HiFi reads have much lower error rates (< 1%), and in comparison with NGS reads are still long, such reads considerably sacrifice on read length in comparison with PacBio CLR and ONT. Overall, short-read genome assembly tools are not directly applicable to long-read data. This explains the need for novel approaches.

Existing methods for haplotype reconstruction of diploid genome from long reads basically fall into two classes. The first one involves alignment based methods, which are referred to as *haplotype assembly* programs. Recent works such as WhatsHap Patterson *et al*. (2015), HapCut2 Edge *et al*. (2017) and HapCol Pirola *et al*. (2016) are designed for diploid haplotype assembly. It is characteristic for these tools to make use of a high-quality reference as a backbone sequence. They align reads to it to call variants using external tools. Subsequently, variants are separated into different haplotypes based on their own phasing algorithms. The output haplotigs of these tools then reflect modified reference sequence patches. However, because only minor modifications of the backbone sequence can be applied, the haplotigs can be affected by non-negligible biases.

For avoiding reference induced biases, *de novo* assembly based methods can be used as an alternative. A series of remarkable *de novo* genome assemblers specialized for long-read sequencing data have been developed so far. FALCON Chin *et al*. (2016), which follows the hierarchical genome assembly process, and its haplotype aware version FALCON-Unzip, are mainly designed for PacBio long reads. Improved Phased Assembler (IPA) PacificBiosciences (2020) is designed to produce phased genome assemblies from PacBio HiFi reads. Canu Koren *et al*. (2017) uses adaptive overlapping algorithm and sparse assembly graph construction which helps to separate repeats and haplotypes, and its modification version HiCanu Nurk *et al*. (2020) is tailored for haplotype aware assembly from PacBio HiFi reads. Moreover, Flye Kolmogorov *et al*. (2019) tries to generate optimal assemblies using repeat graphs. Generic long-read assemblers Shasta Shafin *et al*. (2020) and Wtdbg2 Ruan and Li (2020), considerably speed up the large-scale long-read assembly based on novel graph representations. Notably, Hifiasm Cheng *et al*. (2021) is a haplotype-resolved assembler specifically designed for PacBio HiFi reads. A graph-based approach to haplotype aware diploid assembly was proposed in Garg *et al*. (2018), which combines accurate short-read data and long-read (PacBio) data in a hybrid (employing both NGS and long reads) type framework.

There are various other long read assembly approaches, all of which are not designed to produce haplotype-aware assemblies. They usually collapse homologous sequences into one consensus sequence, and therefore do not distinguish between the haplotypes of a diploid genome sufficiently accurately. We refer the reader to Shafin *et al*. (2020); Logsdon *et al*. (2020) for a comprehensive overview of such tools.

*In summary*, long read assemblers are either designed to work for only specific types of data, such as PacBio HiFi or ONT, or they do not address ploidy by design. In somewhat more detail, there is no method that combines the following three important points: it generates high-quality haplotype-aware genome assemblies only based on long read data (1), is capable of handling all three of PacBio CLR, PacBio HiFi and Nanopore reads (2), and does not depend on high-quality reference sequence as a backbone (3).

Here, we present *phasebook*, and suggest it as an approach that to the best of our knowledge is the first one to address all (1),(2) and (3) in combination. By bridging the gap in the body of existing work, phasebook appears to be the first approach that presents a framework that allows to reconstruct the haplotypes of diploid genomes *de novo* from long reads without having to specialize in a particular sequencing technology.

We evaluate phasebook on different long-read sequencing data, namely PacBio CLR/HiFi reads and Nanopore reads. Benchmarking results on both simulated and real data indicate that our method outperforms state-of-the-art tools (both haplotype assembly and de novo assembly approaches) in terms of various aspects. Thereby, an application scenario of major interest is to reconstruct individual haplotypes of the Major Histocompatibility Complex (MHC) region, spanning ~5 Mbp of chromosome 6, and playing a pivotal role in the adaptive immune system. Because of its great hereditary variability, haplotype identity is of utmost interest with respect to numerous human diseases and the corresponding medical treatments. Overall, phasebook generates more complete, more contiguous and more accurate haplotigs compared with other approaches in this particularly challenging case.

## Results

We have designed and implemented phasebook, a novel approach to assemble individual haplotypes of diploid genomes from long read sequencing data, namely PacBio HiFi, PacBio CLR and Oxford Nanopore reads. In this section, we provide a high-level description of the workflow and evaluate its performance on both simulated and real data, in comparison with an exhaustive selection of existing state-of-the-art tools. These tools include both *de novo* long-read assemblers and reference-based haplotype assembly approaches.

### Approach

See Figure 1 for an illustration of the following description. For full details on the following outline, see descriptions of the individual steps in the workflow figure in Methods.

**Figure 1.**
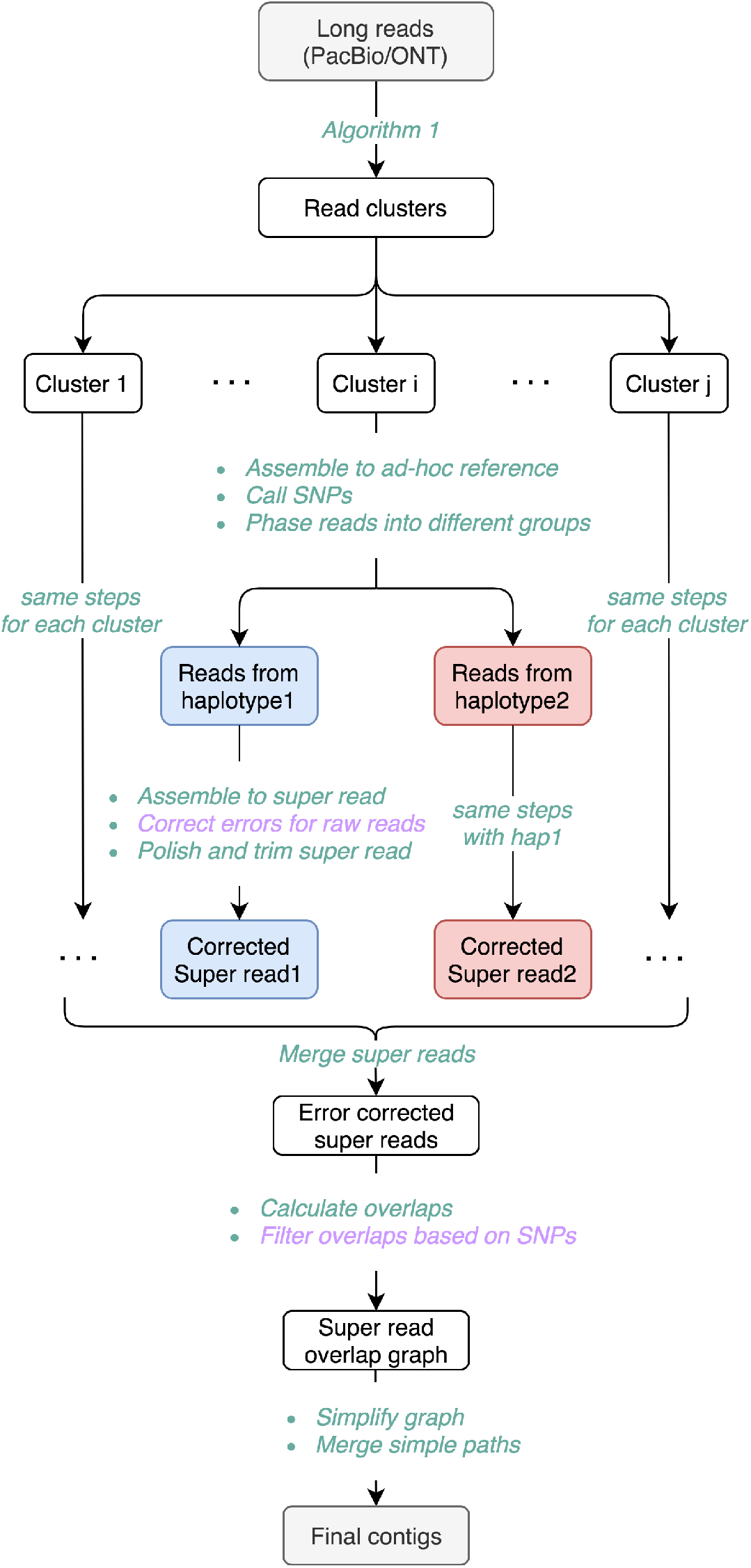
An overview of phasebook. The purple text represents that this step is optional. *Correct errors for raw reads* is recommended for long reads with high sequencing error rate such as PacBio CLR and ONT reads. *Filter overlaps based on SNPs* is recommended for small genomes or specific genomic regions such as MHCs.

#### Workflow: Sketch and Motivation

The approach of phasebook is to first collect contiguous raw reads into small clusters, and compute two haplotype-specific super reads for each of the clusters. Secondly, phasebook constructs an overlap graph where nodes correspond to super reads, and edges indicate that super reads stem from identical haplotypes. Based on the resulting overlap graph, phasebook extends super reads into haplotigs, as the final output.

phasebook is inspired by a few recent approaches, whose virtues are synthesized into an approach that is able to address the universal challenge. First, phasebook is inspired by WhatsHap Patterson *et al*. (2015), as an approach that is based on the minimum error correction (MEC) framework, and excelled in phasing long third-generation sequencing reads in particular, but itself depends on high-quality reference. Here, we design a framework, in which solving the minimum error correction problem can be employed without the need for high-quality reference sequence.

To make this possible, we first determine polymorphic loci using Longshot Edge and Bansal (2019), as a state-of-the-art tool for calling variants in long reads. Then, phasebook constructs an overlap graph on groups of mutually overlapping reads. This allows to align the long reads within the groups with one another. As a result, we can raise a local coordinate systems, which is an essential part of the MEC problem (and which comes without efforts when using reference sequence).

After solving the resulting local MEC instances, phasebook adopts techniques to support haplotype-aware de novo assembly based on overlap graphs, in full consistency with prior steps. In this, phasebook is inspired by recent, related work on short reads that focuses on on overlap graph based assembly (see SAVAGE Baaijens *et al*. (2017) or POLYTE Baaijens and Schönhuth (2019), for example). Here, we adapt these techniques to dealing with long reads. For computing the necessary overlaps, we make use of Minimap2 Li (2018), as the superior current approach to compute overlaps among long reads in sufficiently short time also for large datasets.

#### Workflow: Paradigms and Stages

From a larger perspective, phasebook pursues a divide-and-conquer strategy: in the divide stage, it separates reads into small local groups, which results in many local instances of the *minimum error correction (MEC)* problem. Solving the local MEC instances concludes the divide stage. In the conquer stage, it generates assembly output by following the Overlap-Layout-Consensus (OLC) assembly paradigm. As such, phasebook reflects a combination of an approach that is able to solve the MEC problem without reference sequence, and performing OLC based assembly in a ploidy-aware setting.

#### Workflow: Overview of Technical Steps

Each of the two stages falls into *two basic steps*.

In the *divide* stage, phasebook *first* determines clusters where each of the clusters reflects a collection of reads that overlap each other in terms of genomic position (executed by ‘Algorithm 1’ in Figure 1).

*Secondly, during divide*, phasebook solves an instance of the MEC problem for each such cluster. Thereby, phasebook applies the principles of the original, reference-guided WhatsHap, by implementing them in the de novo setting given here. As above-mentioned, the de novo setting implies the need for an ad-hoc coordinate system as part of the MEC problem. This makes part of methodical challenges that affect this step. As a result, this phases the reads of each such local clusters into two groups. Each such group contains reads that agree in terms of local haplotype. Along with the phasing, it corrects errors in raw reads, and determines an error-corrected super read for each such group. In Figure 1, this comprises all steps from ‘Assemble to ad-hoc reference’ to ‘polish and trim super read’ (where polishing and trimming may be required in low coverage areas). See also Figure 2 for further illustrations. This results in two super reads that reflect local haplotypes for each cluster, one of which generated as per the blue, and one of which generated as per the red branch in Figure 1. We recall that each pair of blue/red super-reads reflects the optimal solution of a local instance of the MEC problem.

**Figure 2.**
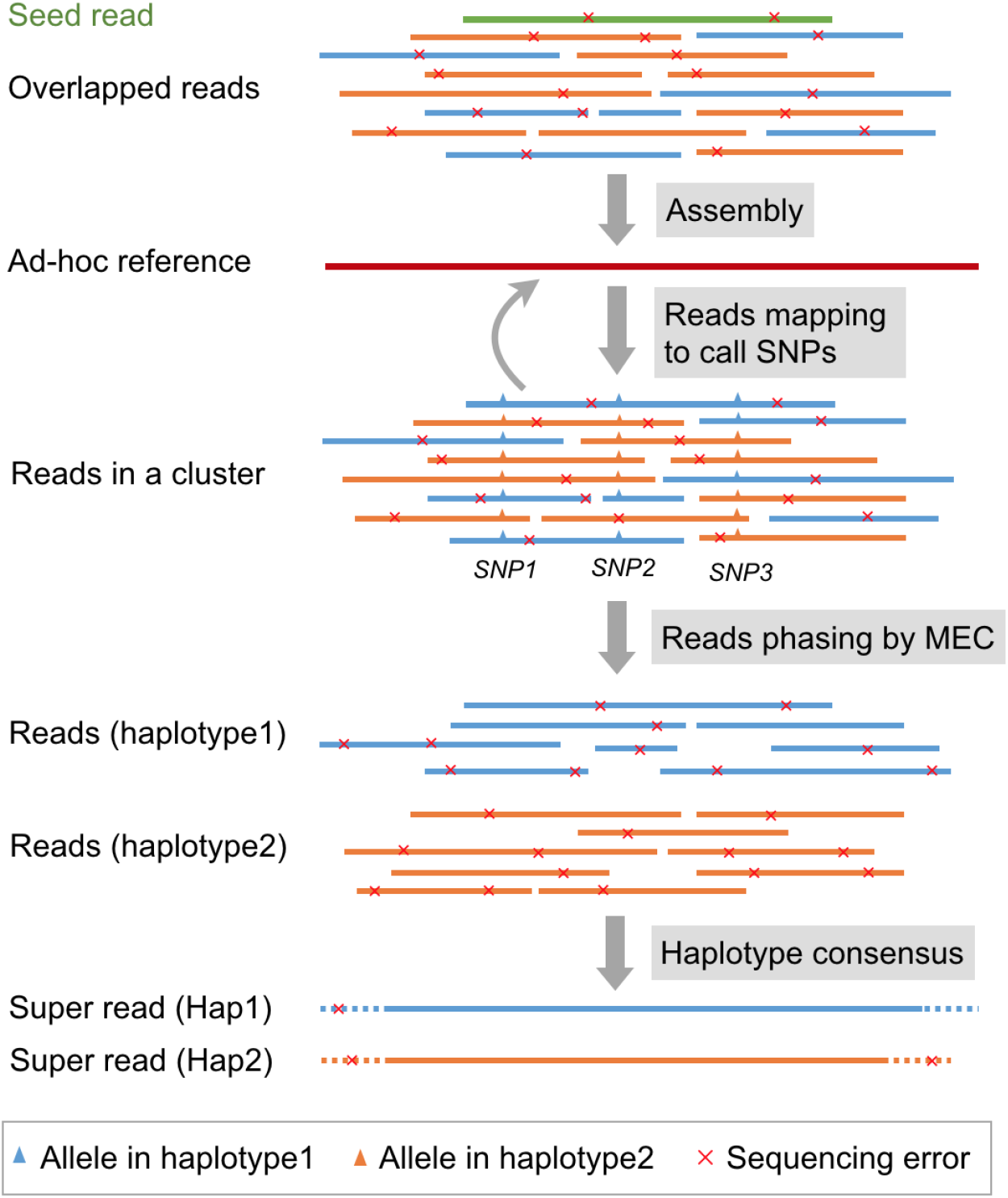
A schematic diagram for reads phasing and super reads generation in a read cluster. The blue reads belong to haplotype 1 and the orange reads belong to haplotype 2. The green read means that this read is a seed read. We use WhatsHap to separate reads into two different groups based on SNPs involved in long reads. The dash line regions in super reads represent the bases with sequencing low coverage, which can be optionally trimmed to generate corrected super reads.

In the *conquer* stage, phasebook *first* constructs an overlap graph on all resulting super reads (comprising all steps from ‘Merge super reads’ to ‘Filter overlaps based on SNPs’ in Figure 1), where edges indicate that two overlapping super-reads stem from the same haplotype, based on sound statistical considerations. Note that the number of super reads is limited which renders the construction of an overlap graph certainly manageable, arguably a virtue of the design of the overall setup. Because super reads are haplotype specific, the corresponding overlap graph virtually consists of two components each of which reflects one of the haplotypes, with only a handful spurious edges connecting the components, see Figure 3 for further illustration.

**Figure 3.**
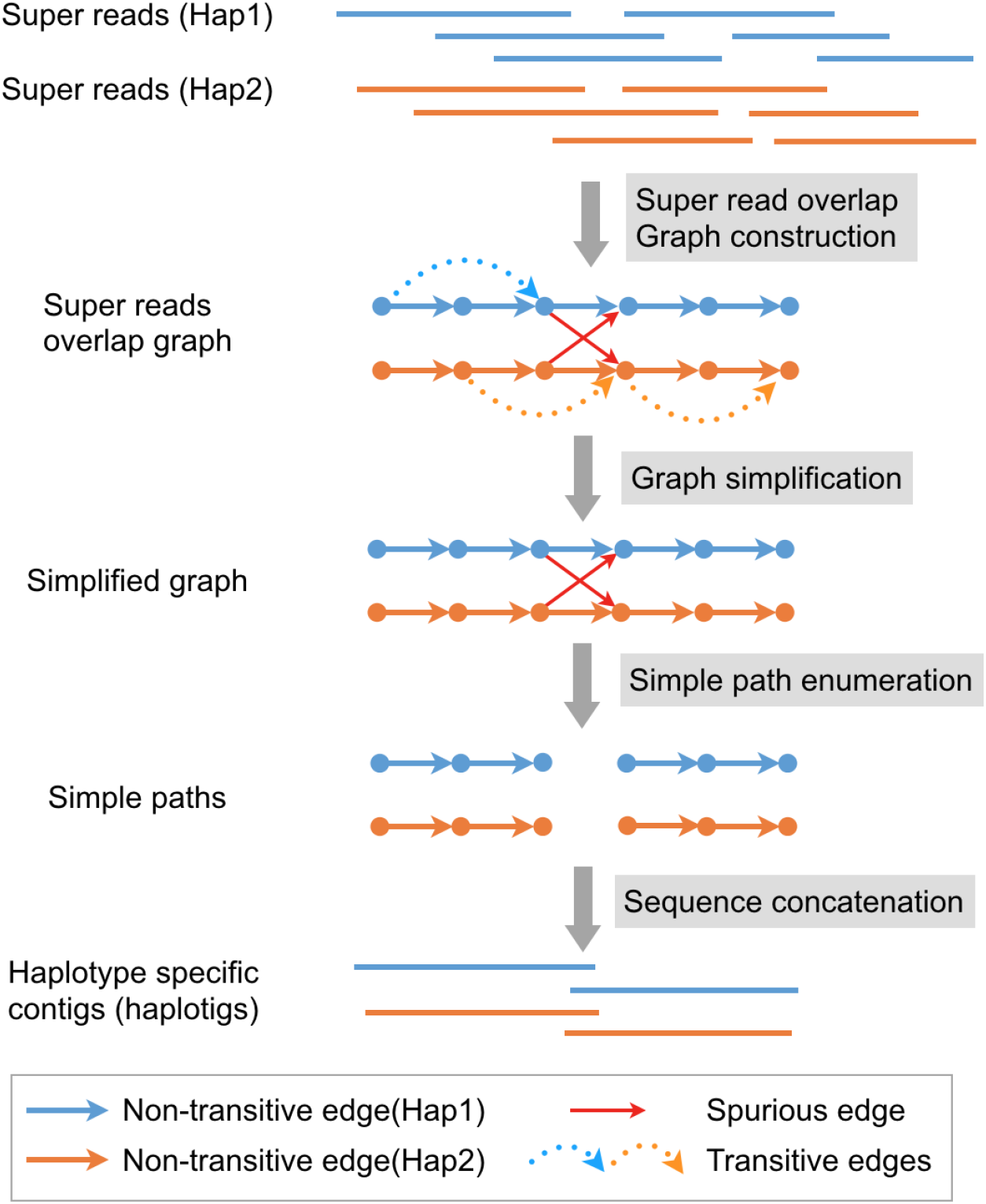
A schematic diagram for super read overlap graph construction. The blue super reads belong to haplotype 1 and the orange super reads belong to haplotype 2. The solid arrow lines (blue or orange) represent non-transitive edges and the dash arrow lines represent transitive edges, respectively. The solid arrow lines (red) represent spurious edges caused by incorrect super read overlaps from different haplotypes.

Finally, in the *second step of the conquer stage*, super reads are assembled into haplotype specific contigs by careful elimination of redundant and spurious edges, where the identification of spurious edges is based on evaluating (putative) polymorphic sites. After edge elimination, phasebook lays out paths in the resulting connected components of the overlap graph, indicated by ‘Simplify graph’ and ‘Merge simple paths’ in Figure 1.

See Methods for full details on each of the sub-steps involved in the overall workflow.

### Datasets

#### Simulated data

We simulated various datasets using different long read sequencing technologies, namely, PacBio CLR, PacBio HiFi and ONT. Generally, PacBio CLR and ONT reads have high sequencing error rate (5% ~15%), while PacBio HiFi reads have much lower error rate (< 1%). We selected two human MHC haplotypes from the Vega Genome Browser: PGF and COX. Subsequently, We used PBSIM Ono *et al*. (2013) model-based simulation, reflecting a sound way to generate PacBio CLR reads of N50 length 20kbp and average sequencing error rate 10% for each of the two haplotypes at a coverage of 25x. We used PBSIM sampling-based simulation to generate PacBio HiFi reads of N50 length 10kbp for each of two haplotypes at a coverage of 15x by utilizing human individual HG00731 as the sample profile, which was downloaded from EBI. Besides, we used NanoSim V2.6.0 Yang *et al*. (2017) as an approved simulator to generate ONT reads of N50 length 20kbp and average sequencing error rate 10% using the built-in pre-trained model `human_NA12878_DNA_FAB49712_albacore` for each of the two haplotypes at a coverage of 25x. For each sequencing platform, the reads from each of the two haplotypes were combined respectively, resulting a read set for a pseudo-diploid genome (PGF and COX). Additionally, we used PBSIM model-based simulation to generate PacBio CLR reads of N50 length 20kbp and average sequencing error rate 10% for each of the two haplotypes at a coverage of 15x, 25x, 35x and 45x, respectively, in order to evaluate the effect of sequencing coverage.

#### Real data

To evaluate our method on real sequencing data, we downloaded PacBio HiFi and CLR sequencing reads of human individual HG00733 from EBI and SRA (accession number: SRR7615963), and Oxford Nanopore PromethION sequencing reads of human individual NA19240 from ENA where the project ID is PRJEB26791. Subsequently, we extracted reads primarily aligned to chromosome 6, which is ~70 Mbp in length. The approximate sequencing coverage per haplotype of the chromosome 6 for these three types of long reads is 15x, 40x and 25x, respectively. Full length haplotypes have been reconstructed for HG00733 and NA19240 in a recent study Chaisson *et al*. (2019) by applying multi-platform sequencing technologies and specialized algorithms. Hence, we downloaded the corresponding haplotype sequences as the ground truth for benchmarking experiments with focus on chromosome 6. More details about data sources are shown in “Availability of data and materials”.

### Run other approaches for comparison

#### Reference-guided haplotype assembly

Haplotype assembly methods such as WhatsHap and HapCut2 take as input a reference genome sequence and a variant set (unphased VCF file), and output a phased VCF file. For simulated data, we selected another MHC haplotype (named DBB) as the reference also from the Vega Genome Browser. For real data, we extracted the sequence of chromosome 6 from GRCh38 as the reference. Subsequently, we aligned sequencing reads to reference genomes and called variants using Minimap2 and Longshot, respectively. Ultimately, we followed the method proposed in Baaijens and Schönhuth (2019) to reconstruct contigs for each haplotype block from phased VCF files using `bcftools consensus`.

#### De novo haplotype aware assembly

For the sake of a fair comparison of the performance of the assembly programs, we chose to use the results which keep the haplotype information rather than collapse it. Consequently, we used the output file `prefix.unitigs.fasta` for Canu, the combination of primary contigs and associated contigs for Falcon, we added the option `--keep-haplotypes` when running Flye, and used the output file `prefix.p_utg.fasta` in Hifiasm for haplotype-aware comparisons. For the remaining assemblers, we just compared with the contigs as output, because of the lack of alternative options. Note that we failed to run Falcon-Unzip on our datasets^1^, because no adequate read overlaps were generated likely due to too low coverage, thus only results of Falcon are reported.

### Assembly performance evaluation

The assembly performance was evaluated by means of several commonly used metrics, routinely reported by QUAST V5.1.0 Mikheenko *et al*. (2018), as a prominent assembly evaluation tool. See Section ‘Metrics’ below for specific explanations. Contigs with length less than 20kbp were filtered before evaluation. For reference-guided haplotype assembly approaches, we generated contigs for each haplotype from phased variants, even if haplotypes share highly similar regions. *De novo* assembly approaches, however, can assemble genomic sequences from regions shared by the different haplotypes only once, which can be interpreted misleadingly. To circumvent the issue, we used the parameters `--ambiguity-usage all` and `--ambiguity-score 0.999` so as to obtain reasonable alignments of contigs with both haplotypes. In addition, for evaluation of large genomes such as chromosome 6, we used the default optimal parameters by adding the flag `--large`.

### Metrics

#### Haplotype coverage (HC)

Haplotype coverage is the percentage of aligned bases in the ground truth haplotypes covered by haplotigs, which is used to measure the completeness of the assembly.

#### N50 and NGA50

N50 is defined as the length for which the collection of all contigs of that length or longer covers at least half the assembly. NGA50 is similar to N50 but can only be calculated when the reference genome is provided. NGA50 only considers the aligned blocks (after breaking contigs at misassembly events and trimming all unaligned nucleotides), which is defined as the length for which the overall size of all aligned blocks of this length or longer equals at least half of the reference haplotypes. Both N50 and NGA50 are used to measure the contiguity of the assembly.

#### Error rate (ER) and N-rate (NR)

The error rate is equal to the sum of mismatch rate and indel rate when mapping the obtained contigs to the reference haplotype sequences. Besides, N-rate is defined as the proportion of ambiguous bases (‘N’s).

#### Misassembled contigs proportion (MC)

If a contig involves at least one misassembly event, it is referred to as misassembled contig, which indicates that left and right flanking sequences align to the true haplotypes with a gap or overlap of more than 1kbp, or align to different strands, or even align to different haplotypes. Here, we report the percentage of misassembled contigs relative to the overall contigs.

### Benchmarking results

We performed benchmarking experiments including all methods on the simulated and real data as described above, for all three types of long reads, namely, PacBio HiFi, PacBio CLR and Oxford Nanopore reads. In summary, the results show that for both simulated and real PacBio HiFi reads, phasebook arguably reconstructs more complete (HC), more continuous (NGA50, except IPA) and more accurate (ER, NR) haplotigs compared with other existing relevant approaches. Advantages of phasebook are most evident for PacBio CLR and Nanopore reads, where our approach outperforms other methods in terms of haplotype coverage on both simulated and real data in terms of a relatively large margin. At the same time, phasebook maintains better or comparable performance in terms of various other metrics (NGA50, ER, NR, and MC). On PacBio HiFi reads, Hifiasm and phasebook share top performance. More details on results are provided in the following sections.

#### PacBio HiFi reads

Table 1 shows the assembly statistics on PacBio HiFi reads. In simulation experiment, phasebook achieves the largest haplotype coverage (HC: 99.4%), while in real data, it reconstructs the second largest part of the haplotypes (HC: 91.2%), rivaled only by Hifiasm (HC: 91.4%), which outperforms phasebook by a relatively small amount. This is arguably compensated by the fact that Hifiasm fails to achieve a large fraction of the haplotype sequence when dealing with the MHC regions (HC: only 11.1%, see ‘Simulated data’). As for assembly contiguity, phasebook achieves the second best NGA50 across all methods except IPA in both experiments while its N50 is much lower than other generic assemblers such as Flye and Wtdbg2, which however is certainly offset that overall assembly length (indicated by HC), to which N50 refers, is small for Flye and Wtdbg2. High N50 is usually further achieved through collapsing small bubbles and smoothing the tangled regions in the assembly graph. This, however, may introduce error bases and misassemblies. An indication of this happening are the high error rate (≥ 10 and 2 ~ 3 times higher than our method in simulated and real data, respectively) and the large proportion of misassembled contigs of Flye and Wtdbg2. HiCanu is the only approach that generates haplotype coverage that is on a par with our method in terms of orders of magnitude (only 1.6% lower) in both datasets, while maintaining the lowest proportion of misassembled contigs (HiCanu: 1.4% vs phasebook: 12.5%, where phasebook comes in clear second however, comparing with all methods supporting sufficient haplotype coverage). One reason for the reduced misassembly rate of HiCanu may be the short length of contigs overall, as indicated by N50 and NGA50 values that are half as large as those of phasebook.

**Table 1.** Benchmarking results for PacBio HiFi reads. HC = Haplotype Coverage, ER = Error Rate (mismatches + indels), NR = N-Rate (ambiguous bases), MC = Misassembled contigs proportion. Top: simulated diploid data (sequencing coverage is 15× per haplotype) for the MHC region. Bottom: real diploid data for chromosome 6 of individual HG00733. Note that we compared with IPA, the official PacBio assembler for HiFi reads instead of Falcon.

|  | HC (%) | N50 | NGA50 | ER (%) | NR(%) | MC(%) |
| --- | --- | --- | --- | --- | --- | --- |
| *Simulated data* |  |  |  |  |  |  |
| phasebook | 99.4 | 501076 | 539378 | 0.017 | 0.0 | 12.5 |
| HiCanu | 97.8 | 283691 | 283691 | 0.009 | 0.0 | 1.4 |
| IPA | 81.9 | 1426746 | 818630 | 0.021 | 0.0 | 5.6 |
| Flye | 56.0 | 827817 | 10861 | 0.169 | 0.0 | 42.9 |
| Wtdbg2 | 49.8 | 3347120 | - | 0.212 | 0.0 | 100.0 |
| Hifiasm | 11.1 | 27223 | - | 0.183 | 0.0 | 0.0 |
| WhatsHap | 59.4 | 257599 | 155079 | 0.240 | 9.0 | 20.0 |
| HapCut2 | 56.5 | 257599 | 155079 | 0.245 | 9.0 | 20.0 |
| *Real data* |  |  |  |  |  |  |
| phasebook | 91.2 | 450480 | 491919 | 0.042 | 0.0 | 3.5 |
| HiCanu | 89.6 | 235876 | 206729 | 0.109 | 0.0 | 2.2 |
| IPA | 73.3 | 11997148 | 722645 | 0.073 | 0.0 | 2.5 |
| Flye | 55.1 | 16247741 | - | 0.084 | 0.0 | 41.2 |
| Wtdbg2 | 54.8 | 33738322 | - | 0.101 | 0.0 | 55.6 |
| Hifiasm | 91.4 | 446213 | 380384 | 0.090 | 0.0 | 4.2 |
| WhatsHap | 76.5 | 381196 | 359841 | 0.058 | 0.4 | 3.3 |
| HapCut2 | 76.5 | 381196 | 354305 | 0.058 | 0.4 | 3.2 |

The currently most predominant reference-guided methods (WhatsHap and HapCut2) only yield less than 60% and 77% haplotype coverage in MHC and Chromosome 6 datasets, respectively. Meanwhile, they have N50 and NGA50 about twice as short, and higher error rates in comparison to our approach. Moreover, reference-guided methods strongly depend on high-quality reference, which is generally hard to generate for highly diverse genomic regions such as the MHC region. Due to the ambiguous bases (‘N’s) in the reference genomes, we therefore observe that the final assemblies generated from reference-guided methods harbor 9.0% and 0.4% N-rate in MHC and Chromosome 6 datasets, respectively. De novo assemblies, however, do not suffer from biases that are inherent to reference genome based approaches, hence do not contain any ‘N’.

#### PacBio CLR reads

The assembly results for PacBio CLR reads are shown in Table 2. phasebook achieves the largest haplotype coverage in both simulated (HC: 95.2%) and real (HC: 92.9%) data compared with all other approaches (HC: 50% ~ 76% throughout). Besides, it outperforms all other de novo approaches in terms of NGA50 (quite dramatically) whereas its N50 is shorter in comparison to others. phasebook also achieves the lowest error rate across all methods in simulated data (MHCs). While other de novo approaches such as Canu and Flye perform better than our method in terms of error rate in real data (Flye ER: 0.079 and Canu ER: 0.124 versus phasebook ER: 0.198), they obtain much lower haplotype coverage (60% and 52%, respectively versus 92,9 from phasebook), higher misassembled contigs proportion, and smaller NGA50 values.

**Table 2.**
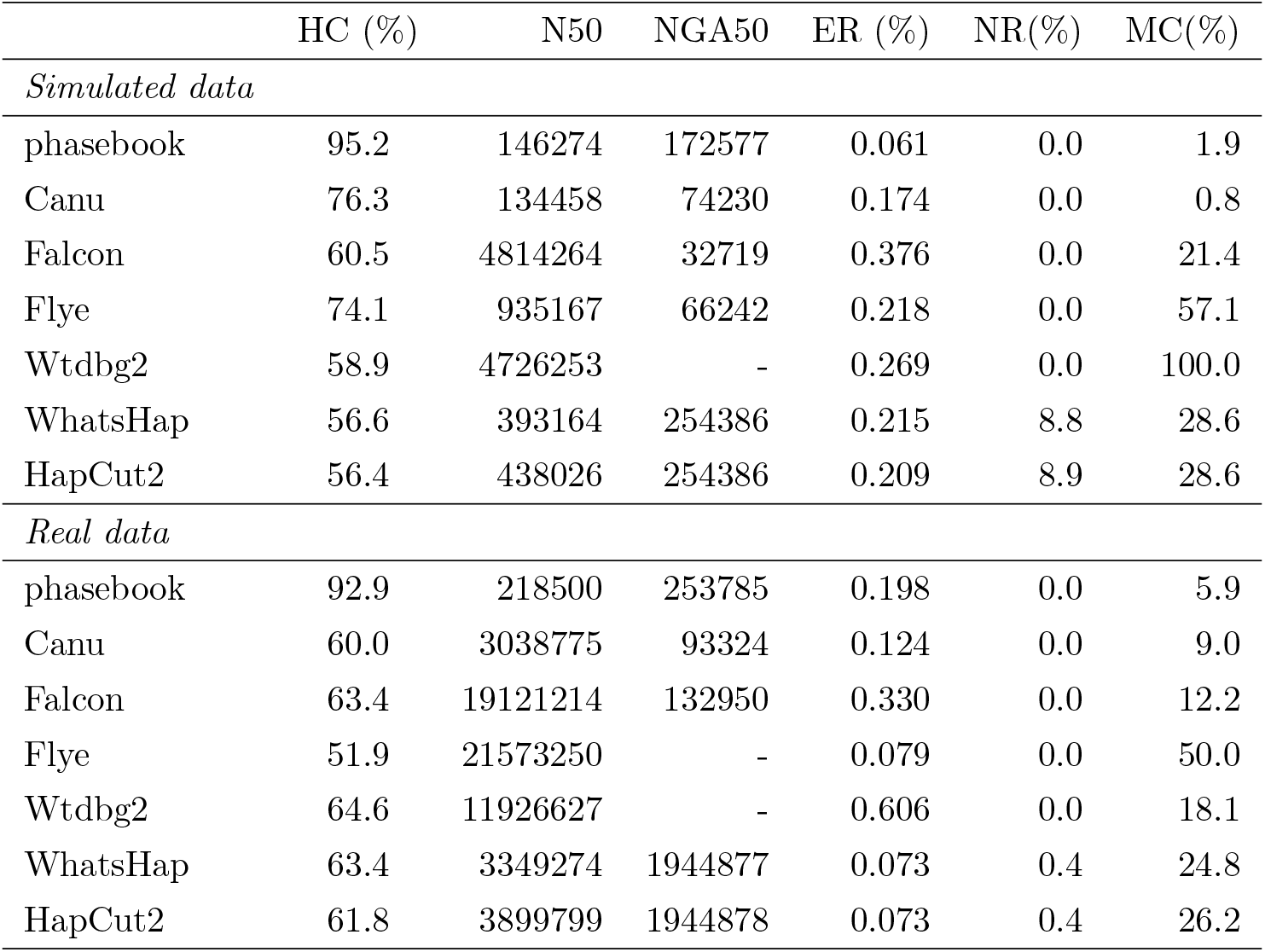
Benchmarking results for PacBio CLR reads. Top: simulated diploid data (sequencing coverage is 25x per haplotype) for the MHC region. Bottom: real diploid data for chromosome 6 of individual HG00733.

Reference-guided methods generate more contiguous haplotigs (larger N50 and NGA50 values). Nevertheless, as we already observed in PacBio HiFi read data, this comes at the cost of substantially elevated N-rates and misassemblies. Additionally, these methods only reconstruct 56% ~ 63% haplotype fraction, which is much lower compared with our method.

#### Oxford Nanopore reads

Table 3 shows the benchmarking results for Oxford Nanopore reads assembly. phasebook still achieves the largest haplotype coverage in both simulated (HC: 97.0%) and real (HC: 84.4%) data compared with all other approaches (HC: 56% ~ 77% in simulated data, 56% ~ 65% in real data). In addition, it generates haplotigs with the lowest error rate (about 2 ~ 11 times lower) in simulated MHC data, and with the lowest misassembled contigs proportion in both simulated and real datasets. Our approach outperforms other de novo assemblers in terms of NGA50 in the real data while its N50 value is still much smaller than others, which is similar to that we already observed in PacBio HiFi/CLR read datasets.

**Table 3.** Benchmarking results for Oxford Nanopore reads. Top: simulated diploid data (sequencing coverage is 25x per haplotype) for the MHC region. Bottom: real diploid data for chromosome 6 of individual NA19240.

|  | HC (%) | N50 | NGA50 | ER (%) | NR(%) | MC(%) |
| --- | --- | --- | --- | --- | --- | --- |
| *Simulated data* |  |  |  |  |  |  |
| phasebook | 97.0 | 108774 | 152599 | 0.092 | 0.0 | 2.8 |
| Canu | 56.8 | 1056827 | 119066 | 0.171 | 0.0 | 17.4 |
| Falcon | 67.8 | 2489921 | 442832 | 0.375 | 0.0 | 20.0 |
| Flye | 54.5 | 1274112 | 86893 | 0.181 | 0.0 | 50.0 |
| Wtdbg2 | 76.5 | 4821843 | 208838 | 0.834 | 0.0 | 33.3 |
| Shasta | 63.0 | 3859518 | 136119 | 1.005 | 0.0 | 33.3 |
| WhatsHap | 56.9 | 493723 | 279218 | 0.204 | 8.9 | 50.0 |
| HapCut2 | 55.9 | 493723 | 279218 | 0.211 | 8.9 | 46.2 |
| *Real data* |  |  |  |  |  |  |
| phasebook | 84.4 | 84379 | 85595 | 1.988 | 0.0 | 9.6 |
| Canu | 65.8 | 4619460 | - | 1.289 | 0.0 | 33.5 |
| Falcon | 64.0 | 4764209 | 37993 | 1.895 | 0.0 | 29.5 |
| Flye | 57.3 | 11457239 | - | 0.686 | 0.0 | 59.3 |
| Wtdbg2 | 60.6 | 15480861 | - | 1.703 | 0.0 | 37.3 |
| Shasta | 62.7 | 2212948 | - | 2.275 | 0.0 | 66.2 |
| WhatsHap | 56.4 | 27552232 | 3520811 | 0.775 | 0.4 | 62.5 |
| HapCut2 | 56.2 | 27552232 | 3285944 | 0.787 | 0.4 | 62.5 |

Reference-guided methods generate more contiguous haplotigs (larger N50 and NGA50 values) in real data. As we have already observed in PacBio HiFi/CLR datasets, this comes at the expense of significantly higher N-rate and misassemblies. Notably, these methods only obtain approximately 56% haplotype coverage, which is much inferior to the result reconstructed from our reference-free approach.

### Effect of sequencing coverage

In order to evaluate the effect of sequencing coverage on noisy long read assembly performance, we used a two MHC haplotypes (PGF and COX) as a template for a diploid genome, and simulated PacBio CLR reads with different sequencing coverage, namely, 15x, 25x, 35x, 45x per haplotype. Subsequently, we ran phasebook and all other tools on these four datasets. The benchmarking results are shown in Table S1 (see Additional File 1??).

As the sequencing coverage varies, phasebook always achieves the best performance in terms of haplotype coverage (95% ~ 97%), clearly outperforming all other tools (52% ~ 76%). In addition, it yields haplotigs with much lower error rate (~ 0.06%, 2 to 6 times lower) at coverages per haplotype of 25x, 35x and 45x. Only at a coverage of 15x, it performs worse than other approaches (phasebook: 0.33% versus 0.217% from Canu, as the best performing de novo assembler). The explanation for this effect is that phasebook corrects sequencing errors at haplotype level, whereas other methods correct errors by considering reads from the two haplotypes together, which yields improvement at low coverage rates, while not being able to improve error rates on increasing coverage. Note as well that it is difficult to correct sequencing errors in noisy long reads when coverage is low, with difficulties arising from 15x and less. In all coverage settings, phasebook computes very little misassembled contigs in comparison with the other approaches; Canu is the only approach that has less misassembled contigs, which however is offset by lower HC, smaller NGA50 values and higher error rates. Besides, the haplotype coverage of phasebook slightly improves on increasing sequencing coverage, and the error rate of phasebook is improved when sequencing coverage increases from 15x to 25x, whereas only slight improvements are observed when raising sequence coverage further to 35x and 45x.

### Runtime and memory usage evaluation

The runtime of phasebook is dominated by two steps. First, the computation of all-vs-all long read overlaps for which we use Minimap2, as one of the fastest approaches to compute read overlaps. Second, generating corrected super reads consumes a considerable amount of time, as a consequence of the high rate of sequencing errors in long reads.

Of note, the number of read clusters is approximately linear with the genome size, as a logical consequence of the design of the clusters. Converting massive amounts of raw reads into a much reduced set of super reads, parallelized across clusters, and only then constructing overlap graphs on (the reduced amount of) super reads, and deriving assemblies from this (comparatively small) graph greatly speeds up the process, and avoids consumption of excessive amounts of memory.

In an overall account, our approach is perfectly apt for assembling large genomes, such as human genomes or other diploid genomes of similar length.

We evaluated the runtime and memory usage of related tools on both simulated and real data (Chr6) for three kinds of long read data, namely, PacBio HiFi, PacBio CLR and Oxford Nanopore reads, and the results are shown in Table S2, S3, S4 (see Additional File 1), respectively. In detail, Table S2 shows that phasebook takes 64.8h CPU time on Chr6 data for PacBio HiFi reads, which is more than other approaches, while not different by orders of magnitude (other de novo approaches: 10. 3 to 41 .0h). In terms of peak memory usage, phasebook requires roughly similar amounts like Canu and Flye. As was to be expected, reference-guided methods, such as WhatsHap and HapCut2, have advantages in terms of runtime and space consumption.

Table S3 shows that phasebook, Canu and Falcon are orders of magnitude slower when assembling CLR reads in comparison with the much more accurate HiFi reads. phasebook requires 1129h CPU time on Chr6 data, which is comparable with Falcon. Table S4 indicates that phasebook takes 727h CPU time on Chr6 data, which is faster than Canu (1139h), but somewhat slower than Falcon (508h). Shasta is the fastest tool for Nanopore reads assembly among all methods we are comparing.

In summary, on real data our approach generally is among the slowest, while, however, never being slower than by a factor of 1 ~ 1.5. In addition, phasebook is well behaved in terms of memory usage. This demonstrates that our approach is very well applicable in all real world scenarios of interest.

## Discussion

Haplotype aware assembly of diploid genomes, which preserves haplotype identity of the assembled sequences, plays a crucial role in various applications. Long-read sequencing data appears to be particularly suitable for addressing this task since long reads can capture linkage of genetic variations across much longer ranges than short reads. However, the elevated error rates of long reads induce considerable difficulties, because it can be challenging to distinguish errors from true variations. The error rates are the major reason why existing long-read assemblers tend to collapse assemblies into consensus sequence, thereby loosing haplotype information. Further, the majority of existing methods for generating haplotype-aware assemblies tends to require guidance by reference genomes. The difference in error profile characteristics of the most popular long-read technologies, such as PacBio HiFi, PacBio CLR and Oxford Nanopore add to the challenge, because it seems that each sequencing technology requires a specially tailored tools, where each tool integrates the particularities of the individual sources of errors. In summary, this explains why prior methods were struggling to address to generate haplotype-aware de novo assemblies for diploid genomes for all predominant sequencing technologies, despite the currently overwhelming interest in such tools.

We have introduced phasebook to overcome this gap in terms of existing methods: phasebook is (to the best of our knowledge) the first method that successfully addresses all of these (crucial) points. phasebook reconstructs the haplotypes of diploid genomes, supporting usage of all predominant sequencing technologies, namely PacBio HiFi, PacBio CLR and Oxford Nanopore, *de novo*, that is without requiring external guidance. Results demonstrate the superiority of the new approach in terms of the majority of approved assembly related metrics.

Key to success for phasebook is to not just follow standard assembly paradigms (such as de-Bruijn or overlap graph based assemblies). Rather, phasebook adopts ideas from the original (non-de-novo) WhatsHap, as a predominant tool for reference-guided read based phasing, and reframes these ideas in a reference-free setting.

In somewhat more detail, phasebook casts the overall assembly problem as several local instances of the minimum error correction (MEC) problem, and assembles the resulting solutions of the local instances thereafter. The challenge inherent to this is to establish the instances of the MEC problem correctly; while this is straightforward in a reference-aided setting, the setup of appropriate MEC instances in a de novo, that is reference-free, setting requires careful consideration of various issues.

Solving the instances corresponds to determining local haplotype-specific patches of genomics sequence. Just as for the original WhatsHap these patches are extremely accurate in terms of error content. This kills two birds with one stone: the local sequence patches do not only account for accurately preserving haplotype identity, but also for correcting errors in the original (long) reads. The high accuracy of the resulting sequence patches, referred to as super-reads, then allows for a fairly straightforward procedure that finalizes the generation of high-quality assemblies: One raises an overlap graph on the super reads, and determines the longest paths in this overlap graph. Because super-reads are 1) long, 2) accurate, and 3) haplotype-specific, the resulting long and accurate overlaps reliably identify the haplotypes also across regions of the genomes that extend the read length by considerable amounts: Full-length assemblies can be easily determined by computing longest paths in the overlap graph. To further polish assemblies, we implement options for sequencing error correction and read overlap refinement that account for differences in terms of sequencing platform characteristics.

Benchmarking results on both simulated and real data indicate that our approach achieves the best performance in terms of haplotype coverage, while maintaining better or comparable performance in terms of various other relevant aspects. Our comparisons consider all state-of-the-art tools that address to assemble diploid genomes in a way that takes ploidy into account. Note that some of the methods are reference-guided, while others generate *de novo* assemblies.

Among the three kinds of long-read sequencing data, PacBio HiFi data is the most promising one in terms of distinguishing the different haplotypes thanks to its low error rates. PacBio CLR and Nanopore reads, on the other hands, come with elevated error rate. As usual, combining long and highly erroneous reads with accurate short reads may establish a viable alternative to achieving even better performance than phasebook, at the expense, of course, of additional efforts in terms of sequencing experiments.

In summary, the results demonstrate that phasebook successfully reconstructs individual haplotypes of the most polymorphic region (MHC) in the human genome at utmost quality in terms of completeness, contiguity and accuracy. The integration of phasebook in population-scale long read sequencing projects may open up novel opportunities.

Still, there is room for improvements. Most importantly, phasebook is confined to work with diploid genomes, so cannot handle polyploid genomes. We consider it worthwhile future work to expand on the idea of polyploids and to design and implement a method that can deal with polyploid genomes, or even cancer and metagenomes, while adopting the ideas of WhatsHap style approaches further.

## Conclusions

We have presented phasebook, a novel approach for reconstructing the haplotypes of diploid genomes from long reads *de novo*, that is without the need for a reference genome. The approach is implemented in an easy-to-use open-source tool https://github.com/phasebook/phasebook. Benchmarking results on both simulated and real data indicate that our method outperforms state-of-the-art tools (both haplotype assembly and de novo assembly approaches) in terms of haplotype coverage, while preserving competitive performance or even achieving advantages in terms of all other aspects relevant for genome assembly. Therefore, the integration of phasebook may reveal new opportunities in population-scale long read sequencing projects.

## Methods

### Read clusters generation

#### Read overlaps calculation

We firstly compute all-vs-all overlaps for raw reads using Minimap2 Li (2018), whose seed-chain-align procedure is known to perform pairwise alignments extremely fast, so can manage the large amount of read pairs we need to process. Secondly, resulting bad overlaps are filtered by reasonable, additional criteria. For example, overlaps that are too short, that do not exceed a minimum level of sequence identity, that reflect self-overlaps, duplicates or internal matches are removed. In this, we follow Algorithm 5 in Li (2016).

#### Read clusters

We sort reads by their length in descending order, considering that longer reads tend to have more overlaps. Processing reads in the corresponding order therefore results in larger read clusters, which reduces the number of clusters, increases the length of the resulting super-reads and hence improves the assembly overall. In each iteration, the longest read having remained unprocessed is selected as the seed read. The corresponding cluster is then determined as the set of reads that overlap the seed read (according to the criteria listed above), and all of its reads are discarded from the sorted list of reads. The procedure stops when all reads have been processed. See the details in Algorithm 1.

##### Algorithm 1 Generate read clusters

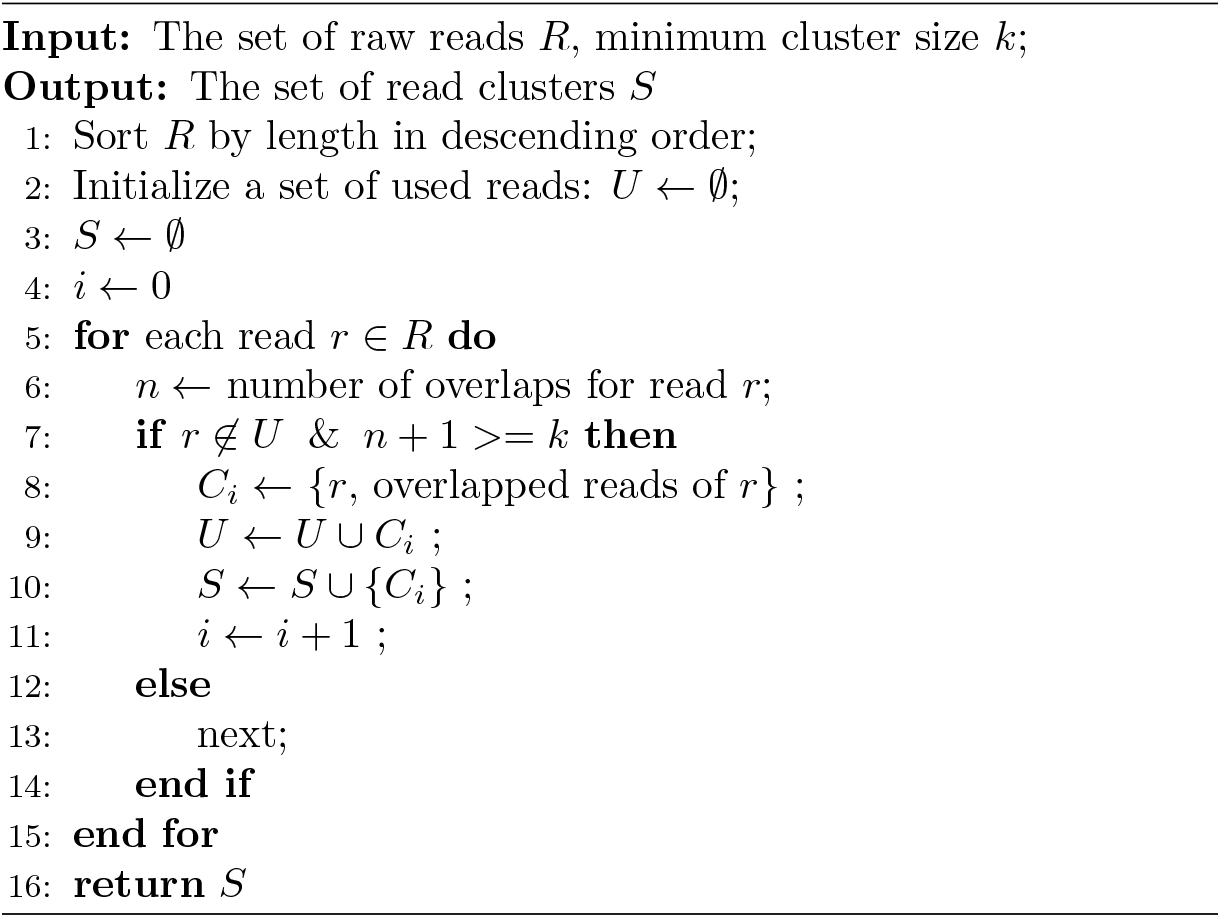

### Reads phasing

Since read clusters contain reads from both haplotypes, we further phase reads into two haplotype-specific groups. See Fig. 2 for the explanation of this step.

#### Read overlap graph

For each read cluster, we largely follow the method proposed in Baaijens *et al*. (2017); Baaijens and Schönhuth (2019) to construct a read overlap graph *G* = (*V, E*) where nodes *v* ∈ *V* represent reads and edges *e* = (*v,w*) ∈ *E* indicate overlaps that are consistent in terms of length, sequence identity (see ‘Parameter settings’). Edges (*v,w*) are directed indicating that the suffix of *v* is likely to be identical with the prefix of *w*. Note that the overlaps were already determined when computing clusters. Appropriate thresholds for sequence identity and length of the overlaps are user defined.

#### Ad-hoc reference construction

In order to reduce the complexity of the overlap graph, we remove transitive edges, which reflects a common procedure. An edge *u* → *v* is called transitive if there is a vertex w and edges *u* → *w,w* → *v*. Subsequently, we determine the longest path in this directed acyclic graph, and the reads that give rise to it. The resulting reads are then assembled into an ad-hoc reference by concatenating the overlapping reads that give rise to the path.

#### SNP calling

To remove remaining sequencing errors, and improve variant (SNP) calling and phasing, we polish the ad-hoc reference using Racon Vaser *et al*. (2017). The raw reads are then mapped back to the ad-hoc reference. SNPs are detected by utilizing a pair-Hidden Markov Model, as implemented in Longshot Edge and Bansal (2019).

#### Reads phasing

Reads within a cluster can stem from both haplotypes. One can phase the reads conveniently, because the majority of them tend to be sufficiently long to bridge neighboring polymorphic sites. For performing the phasing, we make use of WhatsHap Martin *et al*. (2016), as a tool that has been shown to operate at superior performance rates when phasing long reads. WhatsHap centers on the weighted minimum error correction problem, providing a solution with runtime linear in the number of variants. The solution corresponds to partitioning the reads in two groups, each of which immediately corresponds to one of the two local haplotypes of the diploid genome.

### Corrected super reads generation

#### Super read construction

We construct a super-read for each of the two haplotype-specific groups of reads resulting from the phasing procedure. For each read group, we construct another read overlap graph, now comprising only reads as nodes that belong to the same haplotype. Using the overlap graph, we then assemble the reads in each group into a long haplotype specific sequence, which we refer to as *super-read*. As a result, we obtain two super reads for each read cluster (see Fig. 2).

#### Error correction for raw reads

This step is optional and recommended for long reads with high sequencing error rate such as PacBio CLR and Nanopore reads, while it is not necessary when processing PacBio HiFi reads because of their great accuracy. For small genomes or specific genomic regions such as MHCs, we make use of a method for self-correction of long reads in each read group. This method combines a multiple sequence alignment (MSA) strategy with local de Bruijn graphs, and is implemented in CONSENT Morisse *et al*. (2020). When dealing with large genomes (e.g. Chr6), to increase efficiency, we use the MSA-based error correction modules built in MECAT2 Xiao *et al*. (2017) and NECAT Chen *et al*. (2020) for PacBio CLR and Nanopore reads, respectively. Reads in one group are supposed to stem from the same haplotype, thus one can avoid the elimination of true variation, which happens when reads from different haplotypes are mixed together. Therefore, because of the prior arrangement of partitioning reads into haplotype-specific groups, the corrected reads enjoy error rates that are lower than when running these tools on reads referring to unphased settings.

#### Super reads polishing and trimming

Using the error corrected reads, we return to the super-reads computed earlier, and polish each super-read by comparing it with the now error corrected reads. (When using PacBio HiFi reads, we apply the procedure without prior correction of errors.) As a result, the super-reads come with error rates that are comparable to the ones obtained for the error-corrected raw reads. We finally trim the super read, removing parts of low coverage of raw reads. The final result is an error corrected super-read.

Note that the steps in Section ‘Reads phasing’ and Section ‘Corrected super reads generation’ can be parallelized conveniently across clusters, because computations within clusters are independent of other clusters.

### Super read overlap graph construction

#### Super read overlap graph

The super read overlap graph *G*′ = (*V*′, *E*′) is very similar to the read overlap graph, except that it is constructed from the set of corrected super reads instead of raw reads (see Fig. 3). Thus, each node *v* ∈ *V*′ corresponds to a super read and there is an edge *v_i_* → *V_j_* ∈ *E*′ if they have sufficient overlap length and sequence identity. For the last point, consider that the super-reads exhibit very low sequencing error rates, so near-perfect identity in the overlap is a necessary condition for edges.

Because the number of super reads is substantially smaller than the number of raw reads, the super read overlap graph can be constructed in short time. For constructing it, we again compute all-vs-all super read overlaps using Minimap2, filter bad overlaps (remove self-overlaps, internal overlaps, etc.), and eventually establish the overlap graph analogous to the procedure of Section ‘Reads phasing’, where the only difference is that parameters differ; in general, because of the low error rates and the extended length of super reads, more stringent thresholds are necessary.

#### SNP-based spurious edges removal

As there may still be errors in super reads, one still needs to distinguish between uncorrected errors and true variations. As a consequence, super reads can have false positive overlaps when considering sequence identity in the overlapping regions alone, which leads to the introduction of misassemblies.

To control false positive overlaps, we further refine the edges of the super read overlap graph by applying a heterozygous SNP-based procedure to remove spurious edges that we developed. In more detail, for a pair of overlapping super reads, we collect the corrected reads which belong to the read groups that correspond with the super reads, and map the reads back to the sequence formed by concatenating the pair of super reads along their overlap.

Subsequently, we genotype the variants involved in the overlap regions, and remove overlaps that contain at least *k* heterozygous SNPs. In our experiments *k* = 0, which correspond to requiring perfect overlap. This step might be time-consuming if genome size is large. We therefore consider it optional, and recommend it preferably for small genomes or particular genomic regions, such as the MHC region in our case.

### Final contigs generation

#### Graph simplification

First, we simplify the super read overlap graph by removing tips, because tips are likely to be introduced by spurious overlaps, hence reflect mistaken junctures of super reads. Second, we remove all transitive edges, for the exact same reasons for which we remove such edges in the read overlap graph (see Section ‘Reads phasing’ and Fig. 3).

#### Merging simple paths

A node *v* ∈ *V*′ in the directed graph *G*′ is defined as a *non-branching node* if its indegree and outdegree are both equal to one, and *v* is defined as a *branching node* if either its indegree or outdegree is greater than one. A simple path is a maximal non-branching path, that is its internal nodes are non-branching nodes, while starting and final nodes are branching. For such paths, there is only one possible way to combine the corresponding super reads. Therefore, we merge every simple path into a single contig. Since the number of nodes and edges in super read overlap graph *G*′ is small, it is straightforward and very efficient to enumerate and merge all simple paths (see Fig. 3).

#### Final contigs polishing

This step is recommendable when the sequencing coverage is low, i.e. less than 20x per haplotype. The reason is that the ends of super reads are likely to contain an elevated amount of errors. This needs to be addressed and corrected. So, for each contig generated from a simple path, we collect all corrected reads involved in the read groups that gave rise to the super reads forming the simple path, and align them to the contig. By evaluation of the resulting alignment, further errors can be eliminated, resulting in polished contigs as the final output.

### Parameter settings

There are mainly two parameters to be set during the read overlap graph construction, namely, the minimum overlap length and the minimum sequence identity of the read overlap. In our experiments, the minimum overlap length is 1000 bp, and the minimum sequence identity of the read overlap is 0.95 for PacBio HiFi reads, and 0.75 for PacBio CLR and ONT reads, in agreement with the approximate expected amount of identity according to the inherent error rates. These parameters reflect similar choices used with other tools Chin *et al*. (2016); Koren *et al*. (2017).

## Acknowledgements

We would thank Jasmijn Baaijens and Marleen Balvert for their insightful discussions during the early stages of this project. Further, we thank Tobias Marschall for useful comments towards the end of the project, and for suggesting the name of the tool.

## Availability of data and materials

Simulated data can be downloaded from https://doi.org/10.5281/zenodo.4603608. The data used to generate PacBio HiFi sample profile when simulating HiFi reads can be downloaded from EBI http://ftp.1000genomes.ebi.ac.uk/vol1/ftp/data_collections/HGSVC2/working/20190925_PUR_PacBio_HiFi/. For real sequencing data, PacBio HiFi/CLR reads of human individual HG00733 can be downloaded from EBI http://ftp.1000genomes.ebi.ac.uk/vol1/ftp/data_collections/HGSVC2/working/20190925_PUR_PacBio_HiFi/ and SRA (accession number: SRR7615963), respectively. Oxford Nanopore PromethION sequencing reads of human individual NA19240 can be downloaded from ENA (project ID: PRJEB26791). As for ground truth reference sequences used for benchmarking experiments, three human MHC haplotypes (PGF, COX and DBB) are from the Vega Genome Browser http://vega.archive.ensembl.org/info/data/MHC_Homo_sapiens.html. Full length haplotypes (chromosome 6) of individuals HG00733 and NA19240 can be downloaded from ftp://ftp.1000genomes.ebi.ac.uk/vol1/ftp/data_collections/hgsv_sv_discovery/working/20180227_IlluminaPolishedPhasedSV/.

The source code of phasebook is GPL-3.0 licensed, and publicly available at https://github.com/phasebook/phasebook.

## Ethics approval and consent to participate

Not applicable.

## Consent for publication

Not applicable.

## Authors’ contributions

XL and AS developed the method. XL, XK, and AS wrote the manuscript. XL and XK conducted the data analysis. XL implemented the tool. All authors read and approved the final version of the manuscript.

## Footnotes

1 We contacted the authors, but received no response, note that the latest update in the github repository is from 4 years ago. This seems to be in agreement with earlier attempts to contact the falcon-unzip crew Garg *et al*. (2018).

