## Supplementary Material for "phasebook: haplotype-aware de novo assembly of diploid genomes from long reads"

[†]These authors contributed equally to the work.
[*]To whom correspondence should be addressed.

| | HC (%) | N50 | NGA50 | ER (%) | NR(%) | MC(%) |
|---|---|---|---|---|---|---|
| *Coverage 15×* | | | | | | |
| phasebook | 95.7 | 150089 | 159789 | 0.329 | 0.0 | 5.8 |
| Canu | 57.8 | 450976 | 53756 | 0.217 | 0.0 | 5.1 |
| Falcon | 57.1 | 3396726 | 50216 | 0.250 | 0.0 | 11.8 |
| Flye | 53.6 | 946256 | 9724 | 0.221 | 0.0 | 75.0 |
| Wtdbg2 | 52.4 | 2834625 | - | 0.311 | 0.0 | 100.0 |
| WhatsHap | 56.5 | 393164 | 254386 | 0.214 | 8.8 | 37.5 |
| HapCut2 | 56.3 | 393164 | 254386 | 0.211 | 8.9 | 37.5 |
| *Coverage 25×* | | | | | | |
| phasebook | 95.2 | 146274 | 172577 | 0.061 | 0.0 | 1.9 |
| Canu | 76.3 | 134458 | 74230 | 0.174 | 0.0 | 0.8 |
| Falcon | 60.5 | 4814264 | 32719 | 0.376 | 0.0 | 21.4 |
| Flye | 74.1 | 935167 | 66242 | 0.218 | 0.0 | 57.1 |
| Wtdbg2 | 58.9 | 4726253 | - | 0.269 | 0.0 | 100.0 |
| WhatsHap | 56.6 | 393164 | 254386 | 0.215 | 8.8 | 28.6 |
| HapCut2 | 56.4 | 438026 | 254386 | 0.209 | 8.9 | 28.6 |
| *Coverage 35×* | | | | | | |
| phasebook | 97.2 | 81136 | 93110 | 0.056 | 0.0 | 3.4 |
| Canu | 64.3 | 207576 | 95195 | 0.137 | 0.0 | 1.1 |
| Falcon | 64.5 | 3389666 | 120422 | 0.224 | 0.0 | 11.1 |
| Flye | 53.2 | 2707055 | 19559 | 0.254 | 0.0 | 83.3 |
| Wtdbg2 | 70.0 | 4824645 | - | 0.301 | 0.0 | 100.0 |
| WhatsHap | 56.6 | 393164 | 254386 | 0.218 | 8.9 | 35.7 |
| HapCut2 | 56.4 | 438026 | 254386 | 0.213 | 8.9 | 35.7 |
| *Coverage 45×* | | | | | | |
| phasebook | 96.0 | 82206 | 88122 | 0.058 | 0.0 | 6.0 |
| Canu | 62.7 | 211688 | 72489 | 0.113 | 0.0 | 1.2 |
| Falcon | 59.3 | 3408801 | 93028 | 0.225 | 0.0 | 15.0 |
| Flye | 61.7 | 2686109 | 22165 | 0.207 | 0.0 | 80.0 |
| Wtdbg2 | 51.6 | 3454287 | - | 0.247 | 0.0 | 100.0 |
| WhatsHap | 57.4 | 393164 | 254385 | 0.208 | 8.8 | 35.7 |
| HapCut2 | 56.4 | 438026 | 254386 | 0.216 | 8.9 | 35.7 |

**Table S1.** Benchmarking results for PacBio CLR reads with different sequencing coverage. HC = Haplotype Coverage, ER = Error Rate (mismatches + indels), NR = N-Rate (ambiguous bases), MC = Misassembled contigs proportion. Simulated diploid data (sequencing coverage per haplotype is 15x, 25x, 35x and 45x, respectively) for the MHC region.

| | CPU time (h) | peak memory usage (GB) |
|---|---|---|
| *Simulated data* | | |
| phasebook | 1.1 | 4.1 |
| Canu | 0.5 | 5.5 |
| IPA | 0.4 | 0.6 |
| Flye | 1.3 | 2.4 |
| Wtdbg2 | 0.3 | 0.5 |
| Hifiasm | 0.003 | 0.3 |
| WhatsHap | 0.1 | 0.8 |
| HapCut2 | 0.4 | 0.9 |
| *Real data* | | |
| phasebook | 64.8 | 27.3 |
| Canu | 24.3 | 26.8 |
| IPA | 14.4 | 16.9 |
| Flye | 41.0 | 33.2 |
| Wtdbg2 | 22.8 | 8.1 |
| Hifiasm | 10.3 | 7.8 |
| WhatsHap | 2.2 | 6.0 |
| HapCut2 | 2.9 | 6.0 |

**Table S2.** Runtime and memory usage for PacBio HiFi reads. Top: simulated diploid data (sequencing coverage per haplotype is 15x) for the MHC region. Bottom: real diploid data for chromosome 6 of individual HG00733.

| | CPU time (h) | peak memory usage (GB) |
|---|---|---|
| *Simulated data* | | |
| phasebook | 22.0 | 35.3 |
| Canu | 9.1 | 6.1 |
| Falcon | 3.9 | 10.4 |
| Flye | 1.7 | 5.6 |
| Wtdbg2 | 0.6 | 0.6 |
| WhatsHap | 0.2 | 1.8 |
| HapCut2 | 0.5 | 1.8 |
| *Real data* | | |
| phasebook | 1129.2 | 41.8 |
| Canu | 731.2 | 28.4 |
| Falcon | 1006.0 | 23.6 |
| Flye | 103.7 | 46.9 |
| Wtdbg2 | 22.7 | 13.5 |
| WhatsHap | 7.6 | 7.7 |
| HapCut2 | 7.8 | 7.6 |

**Table S3.** Runtime and memory usage for PacBio CLR reads.Top: simulated diploid data (sequencing coverage per haplotype is 25x) for the MHC region. Bottom: real diploid data for chromosome 6 of individual HG00733.

| | CPU time (h) | peak memory usage (GB) |
|---|---|---|
| *Simulated data* | | |
| phasebook | 48.4 | 5.2 |
| Canu | 52.2 | 6.4 |
| Falcon | 6.3 | 10.1 |
| Flye | 1.0 | 5.5 |
| Wtdbg2 | 0.3 | 0.8 |
| Shasta | 0.2 | 1.1 |
| WhatsHap | 0.2 | 1.6 |
| HapCut2 | 0.6 | 1.7 |
| *Real data* | | |
| phasebook | 726.6 | 47.1 |
| Canu | 1138.6 | 16.0 |
| Falcon | 507.6 | 23.6 |
| Flye | 66.1 | 45.5 |
| Wtdbg2 | 25.5 | 11.4 |
| Shasta | 4.7 | 29.9 |
| WhatsHap | 12.5 | 23.4 |
| HapCut2 | 165.2 | 23.4 |

**Table S4.** Runtime and memory usage for Nanopore reads.Top: simulated diploid data (sequencing coverage per haplotype is 25x) for the MHC region. Bottom: real diploid data for chromosome 6 of individual NA19240.
